# Optogenetic stimulation of long-range inputs and functional characterization of connectivity in patch-clamp recordings in mouse brain slices

**DOI:** 10.1101/491563

**Authors:** Louis Richevaux, Louise Schenberg, Mathieu Beraneck, Desdemona Fricker

## Abstract

Knowledge of cell type specific synaptic connectivity is a crucial prerequisite for understanding brain wide neuronal circuits. The functional investigation of long-range connections requires targeted recordings of single neurons combined with the specific stimulation of identified distant inputs. This is often difficult to achieve with conventional, electrical stimulation techniques, because axons from converging upstream brain areas may intermingle in the target region. The stereotaxic targeting of a specific brain region for virus-mediated expression of light sensitive ion channels allows to selectively stimulate axons coming from that region with light. Intracerebral stereotaxic injections can be used in well-delimited structures, such as the anterodorsal thalamic nuclei, and also in other subcortical or cortical areas throughout the brain.

Here we describe a set of techniques for precise stereotaxic injection of viral vectors expressing channelrhodopsin in the anterodorsal thalamus, followed by photostimulation of their axon terminals in hippocampal slices. In combination with whole-cell patch clamp recording from a postsynaptically connected presubicular neuron, photostimulation of thalamic axons allows the detection of functional synaptic connections, their pharmacological characterization, and the evaluation of their strength in the brain slice preparation. We demonstrate that axons originating in the anterodorsal thalamus ramify densely in presubicular layers 1 and 3. The photostimulation of Chronos expressing thalamic axon terminals in presubiculum initiates short latency postsynaptic responses in a presubicular layer3 neuron, indicating a monosynaptic connection. In addition, biocytin filling of the recorded neuron and posthoc revelation confirms the layer localization and pyramidal morphology of the postsynaptic neuron. Taken together, the optogenetic stimulation of long-range inputs in *ex vivo* brain slices is a useful method to determine the cell-type specific functional connectivity from distant brain regions.

## INTRODUCTION

Defining connectivity between brain regions is necessary to understand neural circuits. Classical anatomical tracing methods allow establishing interregional connectivity, and lesion studies help to understand the hierarchical organization of information flow. For example, brain circuits for spatial orientation and head direction signaling involve the directional flow of information from the thalamus to the presubiculum. This has been demonstrated by lesion studies of antero-dorsal thalamic nuclei (ADN) that degrade the head direction signal in the downstream dorsal presubiculum, as well as the parahippocampal grid cell signal ^1,2^.

The functional connectivity between brain areas is more difficult to establish at a cellular and subcellular level. In the hippocampus, a highly organized anatomy allows to investigate pathway specific synaptic connections using electrical simulation in the slice preparation. Stimulation electrodes placed in stratum radiatum of CA1 can be used to specifically stimulate Schaffer collateral input from CA3 ^3^. Stimulating electrodes placed in stratum lacunosum moleculare of CA1 will activate the perforant path input to CA1 in isolation ^4^. Electrical stimulation activates neurotransmitter release from axon terminals, however, it activates neurons with somata near the stimulation site as well as axons of passage. It is therefore of limited use for studying afferents from defined brain regions when fibers of different regions of origin intermingle in the target structure, as is typically the case in the neocortex.

Neurons may also be stimulated with light. Optical methods include the photoactivation of caged glutamate, which can be combined with one or two photon laser scanning: Multiple closely spaced sites may be stimulated sequentially, with no mechanical damage to the tissue ^5^. This has been successfully used to map synaptic receptors or to activate individual neurons ^6^. While glutamate uncaging can be used for local circuit analysis, long-range inputs cannot be activated specifically.

A method of choice for the investigation of the long-range connectivity of neuronal circuits is the use of virus mediated channelrhodopsin expression. Using *in vivo* stereotaxic injections as described here, the expression of light-gated ion channels can be targeted and spatially restricted to a desired brain region. In this way, channelrhodopsins are effective for mapping excitatory or inhibitory connectivity from one region to its target. Transfected axons terminals may be stimulated with light in a brain slice preparation, and patch clamp recordings as a read-out allow to examine the function and strength of specific circuit components in the brain ^7^. The optogentic approach combined with stereotaxic injection of virus offers unprecedented spatial and genetic control ^8^. Stimulating with light allows high temporal and spatial precision ^9,10^.

The presubiculum is a 6-layered cortical structure at the transition of the hippocampus and the para-hippocampal formation ^11,12^. It receives important synaptic input from the ADN ^10^, but also from several other cortical and subcortical regions ^13^. Thus, it is impossible to selectively stimulate thalamic axon terminals in a presubicular slice with electrical stimulation or glutamate uncaging. Here, we describe how to determine functional connectivity between distant brain regions (ADN and presubiculum) using precise stereotaxic injections of viral vectors expressing light-gated channels. We show results from the photostimulation of axons terminals of projecting neurons in their target region, coupled with whole cell patch-clamp recordings of post-synaptic neurons in the brain slice preparation.

## MATERIALS and METHODS

### 1. Planning of the experiment

1.1. Define the brain area to be targeted. Determine the stereotaxic coordinates of the injection site with the help of a mouse brain atlas ^14^. For the right antero-dorsal thalamic nucleus (ADN), the coordinates are −0.82 posterior, 0.75 lateral, −3.2 depth in mm relative to bregma. Confirm and document the exactitude of the coordinates by injecting a fluorescent tracer (Fluororuby) in a pilot experiment (cf. Fig. 1A, B).

**Figure 1.**
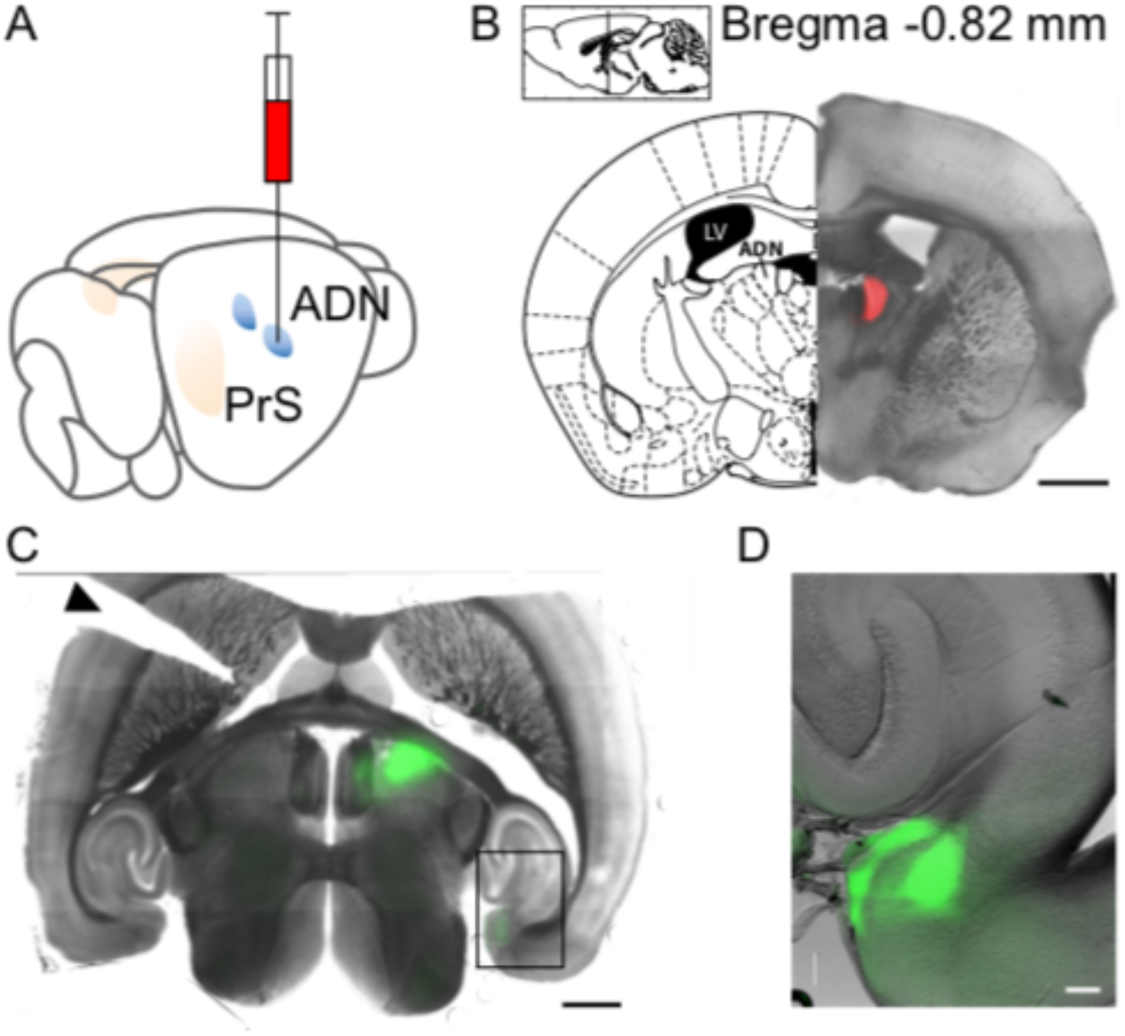
Stereotaxic injection in the anterodorsal thalamic nucleus (ADN). **A,** Schematic representation of injection. **B,** Injection site confirmation with Fluororuby in coronal section. Inset indicates antero-dorsal level and distance from bregma. **C**, Horizontal slice following AAV-Chronos-GFP injection in the thalamus. Note axonal projections to the ipsilateral presubiculum. An incision on the left side of the slice (indicated by a black triangle) marks the contralateral hemisphere. **B-C**, Scale bar 1 mm. **D,** Magnified view of inset in **C** with ADN projections to the presubicular superficial layers 1 and 3. Scale bar, 100 µm.
1.2. Define the type of virus to be injected. Here we use a AAV2/5 serotype expressing Chronos, a fast channelrhodospin-2 variant, fused to green fluorescent protein under the control of the Synapsin promoter: AAV5.Syn.Chronos-GFP.WPRE.bGH (Penn Vector Core, Addgene 59170P). Store the virus in 6 µl aliquots at −80°C as recommended by the producer. Bring 1 aliquot placed on ice to the surgery room, for injecting 1 to 6 animals on a given day. All procedures were performed in accordance with the European Community Council Directive (2010/63/EU) and French Law (87/848).

### 2. Stereotaxic surgery

2.1. Install the stereotaxic frame, and the Hamilton syringe with needle.
2.2. Install the pump. Test the ejection system with water.
2.3. Anesthetize a 4 to 5-week-old C57BL6 mouse with an intraperitoneal injection of a mix of ketamine hydrochloride and xylazine (100 and 15 mg/kg respectively; cf. Table 1). Prepare mix of 1 ml ketamine and 0.5 ml xylazine in 8.5 ml 0.9% NaCl. Inject i.p. 10 µl per gram of the animal’s body weight. Duration of anesthesia is about 1h.

**Table 1.** List of Materials.
2.4. Verify that the animal is well anesthetized with a toe pinch. Then pull out the tongue to facilitate breathing. Shave the cranial hair. To expose the skull, make a straight cut in the scalp with small surgery scissors. Place the animal in a stereotaxic frame, inserting ear bars rostral to the ear canals, and pulling down the skin. Install the nose piece.
2.5. Maintain the body of the animal horizontally at the level of the head using a height-adjusted support. Place a heating pad under the mouse to keep it at physiological temperature.
2.6. Clean the skull by applying 0.9% NaCl with a q-tip to remove soft tissue from the bone.
2.7. Use binoculars for the rest of the surgery.
2.8. Adjust the skull so that the bregma-lambda axis is horizontal, moving up or down the nose and teeth piece. This necessitates iterative measures of bregma and lamba, as both will change following the adjustment of the nose level.
2.9. Find the location of the injection site on the skull. Adjust the injection needle above the injection site according to posterior and medial coordinates and mark the skull with a disposable needle.
2.10. Use a 0.5 mm bur with a drill to realize a 1 mm diameter craniotomy on the mark. Swab eventual bleeding with a paper tissue.
2.11. Fill a 10 µl Hamilton syringe equipped with a 33 gauge needle with a drop of the selected virus disposed on a parafilm. The drop volume should be slightly superior to the desired injection volume (600 nl for 200 nl injected). This will give a safety margin in case some of the liquid is lost during the transfer and allows to produce a small test ejection before proceeding.
2.12. Make sure the syringe has been filled correctly. Verify the functioning of the ejection system by driving down the plunger to test eject a small drop of liquid of 50 nl under visual control. Wipe the drop.
2.13. Down the needle into the brain to the chosen depth. A small volume (50-300 nl, depending on the virus used) is slowly ejected over 10 minutes with an automatic pump.
2.14. Wait 10 min after the injection to avoid leaking from the injection site. Then slowly remove the needle over 3 to 5 minutes.
2.15. Rotate the vertical part of the stereotaxic frame with the syringe away from the animal. Immediately wash the needle in clean distilled water by filling-emptying it several times, in order to avoid clogging. Store the syringe filled with water.
2.16. Remove the mouse from the stereotaxic frame. Suture the skin with 4-0 polyamide suture filament.
2.17. Place the mouse in a heated cage until complete wake up from anesthesia and provide water and food. Ketoprofen (2-5 mg/kg, subcutaneously, 24h) or buprenorphine (0.05-0.1 mg/kg, subcutaneously, 8-12 hours) may be administered to prevent pain.
2.18. Return the animal to its home cage and monitor its well-being. Depending on the virus used, the time for full expression may vary. Here, we allow 3 weeks for expression of AAV5.Syn.Chronos-GFP.

|  | Company | Catalog Number | Comments/Description |
| --- | --- | --- | --- |
| Ketamine 1000 | Virbac |  | dilute 1 mL in 10 mL |
| Rompun 2% (xylazine) | Bayer |  | dilute 0.5 mL in 10 mL |
| Sodium chloride 0.9% | Virbac |  | dilute 8.5 mL in 10 mL |
| Heparin choay | Sanofi | | 100 μL |
| Surgical instruments | FST |  | Scissors, pliers, needle holder, spatula, etc... |
| Stereotaxic frame with digital display | Kopf | Model 940 | Small animal stereotaxic instrument |
| High speed rotary micromotor kit | Foredom | K.1070 |  |
| 0.5 mm bur | Harvard Apparatus | 724962 |  |
| Micro temperature Controller | Physitemp | MTC-1 |  |
| Heating plate | Physitemp | HP4M |  |
| Syringe pump | kdScientific | Legato 130 - 788130 | Infuse and Withdraw modes |
| 10 μL Hamilton syringe | Hamilton | 1701 RN - 7653-01 | |
| Beveled metal needle | Hamilton | 7803-05 | small hub removable needle, 33 gauge, 13mm, point style 4-20° |
| Suture filament Ethilon II 4-0 polyamid | Ethicon | F3210 |  |
| Versi-dry bench absorbant paper | Nalgene |  |  |
| Butterfly needle | Braun | Venofix A | for perfusion |
| Tissue slicer | Leica | VT1200S | cutting parameters : speed 0.07, amplitude 1. |
| Double edge stainless steel razor blades | Electron Microscopy Sciences | 72000 | Use half of the blade in the slicer |
| Filter paper | Whatman |  |  |
| Dual Fluorescent Protein Flashlight | Nightsea | DFP-1 | Royal Blue and Green excitation |
| Upright microscope | Olympus | BX51W1 |  |
| CCD Camera | Andor | DL-604M |  |
| Bath temperature controller | Luigs & Neumann | SM7 | Set at 34°C |
| Peristaltic pump | Gilson | Minipuls 3 | 14-16 ml/min |
| Tubing | Gilson | F117942, F117946 | Yellow/Black, Purple/Black |
| Borosilicate Capillaries | Havard Apparatus | GC150-10 | 1.5 mm outer, 0.86 inner diameter |
| Brown Flaming | Sutter | P-87 |  |
| electrode puller | Instruments |  |  |
| Manipulators | Luigs & Neumann | SM-7 |  |
| Digidata 1440A | Axon Instruments |  |  |
| MultiClamp 700B | Axon Instruments |  |  |
| pClamp software | Axon Instruments |  |  |
| Isolated Pulse Stimulator | A-M Systems | 2100 |  |
| LED Power Supply | Cairn Research | OptoLED Light |  |
| 470 nm LED | Cairn Research |  |  |
| All chemicals | Sigma |  |  |
| 36% PFA | Sigma-Aldrich | F8775 |  |
| 10X PBS solution | Thermofisher | AM9624 |  |
| BupH Phosphate Buffered Saline pack | Thermofisher | 28372 |  |
| Streptavidin-Cy5 conjugate | Thermofisher | S32357 |  |
| Streptavidin-Cy3 conjugate | Life technologies | 434315 |  |
| ProLong Gold antifade mounting medium | Thermofisher | P36390 |  |
| DAPI | Sigma | D9542 |  |
| Milk powder | Carnation |  |  |

### 3. Solutions for acute slice recordings and for fixation

3.1. Prepare stock solutions of 10 times concentrated cutting solution (in mM: 125 NaCl, 25 sucrose, 2.5 KCl, 25 NaHCO_3_, 1.25 NaH_2_PO_4_ and 2.5 D-glucose) and artificial cerebrospinal fluid (ACSF) solution (in mM: 124 NaCl, 2.5 KCl, 26 NaHCO_3_, 1 NaH_2_PO_4_ and 11 D-glucose) in pure deionized water prior to electrophysiology experiments. These solutions are stored at 4°C in 1l bottles without CaCl_2_ and MgCl_2_.
3.2. On the day of the experiment, dilute the stock solutions 10 times to a final volume of 0.5 liter each. Oxygenize by bubbling with 95/5% O_2_/CO_2_. Add divalent ions to obtain final concentrations of 0.1 CaCl_2_ and 7 MgCl_2_ for the cutting solution, and 2 CaCl_2_ and 2 MgCl_2_ for ACSF.
3.3. The potassium-gluconate based pipette solution contains (in mM): 135 K-gluconate, 1.2 KCl, 10 HEPES, 0.2 EGTA, 2 MgCl_2_, 4 MgATP, 0.4 Tris-GTP, 10 Na_2_-phosphocreatine, and 2.7–7.1 biocytin for post-hoc cell morphology revelation. Solution pH is adjusted to 7.3, and osmolarity to 290 mOsm. 1 ml aliquots are stored at −20°C.
3.4. 0.1 M PBS is prepared by diluting BupH Phosphate Buffered Saline dry-blend powder pouches in 500 ml of distilled water, resulting in 0.1 M sodium phosphate, 0.15 M NaCl, pH 7.2.
3.5. To prepare 1 l of 4% PFA solution, dilute 111 ml of 36% liquid PFA and 90 ml of 10X PBS solution in distilled water.
3.6. 30% sucrose solution contains 150 g of sucrose in 500 ml of 0.1 M PBS.

### 4. Preparation of brain slices

4.1. Prepare the bench space with absorbent bench paper for perfusing the mouse via the heart.
4.2. A drip is installed about 1 m above the bench for gravity-fed perfusion. Attach a butterfly needle size 24 G.
4.3. Surround the cutting chamber with ice and store it in a freezer.
4.4. Anesthetize the mouse with intraperitoneal injection of the same ketamine-xylazine mixture as used for surgery. When fully asleep, inject 100 µl of heparin intraperitoneally.
4.5. Fix the animal with adhesive tape on the absorbent paper. Open the thoracic cage, clamp the descending aorta, and perfuse via the left ventricle of the heart with 4°C cooled and oxygenated (95/5% O_2_/CO_2_) cutting solution.
4.6. After 5 seconds, open the right atrium with small scissors.
4.7. After 5 minutes of perfusion, when the organs are bloodless, stop the perfusion, decapitate the animal and immerge the head into cooled and oxygenated cutting solution in a petri dish.
4.8. To extract the brain, cut the skin from neck to nose, section the last vertebra from the skull. Retract the skin and use small scissors to open the skull, cutting it along the midline, from caudal to rostral, up to between the eyes. Carefully remove parietal bone and caudal part of frontal bone. Extract the brain with a small spatula by inserting the instrument between the brain and the cranial floor, sectioning the olfactory bulb, the optic nerve and other cranial nerves, and the cerebellum.
4.9. Gently submerge the brain in ice-cold cutting solution in a beaker.
4.10. Position the brain on filter paper to gently dry the cortical surface. Glue it upside-down to the specimen holder of a vibratome, caudal part facing the blade, in order to cut horizontal brain slices.
4.11. Fill the cutting chamber with ice-cold oxygenated cutting solution so the brain is fully immerged. Make a cut on the left hemisphere (contralateral to the injected side) to avoid potential left-right ambiguity on slices. Caution: always oxygenate the solution and protect slices from light exposure.
4.12. Cut 300 µm-thick slices with the vibratome, at a speed of 0.07 mm/s and 1 mm amplitude.
4.13. At this stage one may briefly check the Chronos-GFP expression in the thalamus using a Nightsea fluorescent lamp and corresponding filter glasses.
4.14. Isolate the hippocampal region with a scalpel and transfer it to a chamber positioned in a beaker filled with bath-warmed (34°C), oxygenated (95/5% O_2_/CO_2_) ACSF.
4.15. After 15 min, take the chamber out of the heated water bath and let the slices rest at room temperature, still oxygenated for at least 45 min and until use.

### 5. Whole-cell patch-clamp recordings

5.1. Gently transfer a brain slice containing the hippocampal complex with a custom made glass transfer pipette to the recording chamber mounted on an upright microscope. The recording chamber (volume 3 ml) is continuously perfused with 34°C warm ACSF bubbled with 95/5% O_2_/CO_2_. The speed of the peristaltic pump is set to 14-16 ml/min.
5.2. Chronos-GFP expression in axon terminals in the region of interest is examined briefly with blue LED illumination and observed with a 4x objective, with a CCD camera image viewed on a computer display.
5.3. Place a slice anchor made from a U-shaped platinum wire with tightly spaced nylon strings (“harp”) on the slice to maintain it.
5.4. Change to a 63x immersion objective and adjust the focus. Check for axons expressing Chronos-GFP and choose a pyramidal neuron for patch recording.
5.5. Move the objective up.
5.6. Pull pipettes using a Brown-Flaming electrode puller from borosilicate glass of external diameter 1.5 mm. Fill pipettes with K-gluconate based internal solution.
5.7. Mount the pipette in the pipette holder on the headstage. Lower the pipette in the chamber, find the tip under the objective. Pipette resistance should be between 3 and 8 MΩ. Apply a light positive pressure with a syringe so as to see a cone of solution outflow out of the pipette and progressively lower the pipette and the objective to the surface of the slice.
5.8. Patch the cell in voltage-clamp configuration: Approach the identified neuron and delicately press the pipette tip onto the soma. The positive pressure should produce a dimple on the membrane surface. Release the pressure to create a giga-ohm seal (>1 GΩ resistance). Once sealed, apply a −65 mV holding voltage. Break the membrane with a sharp pulse of negative pressure.
5.9. Record in current clamp mode the responses of the neuron to hyperpolarizing and depolarizing current steps. This protocol will be used to determine active and passive intrinsic properties of the cell (Fig. 2A). Custom-written Matlab routines are used for off-line analysis ^9,15^.

**Figure 2.**
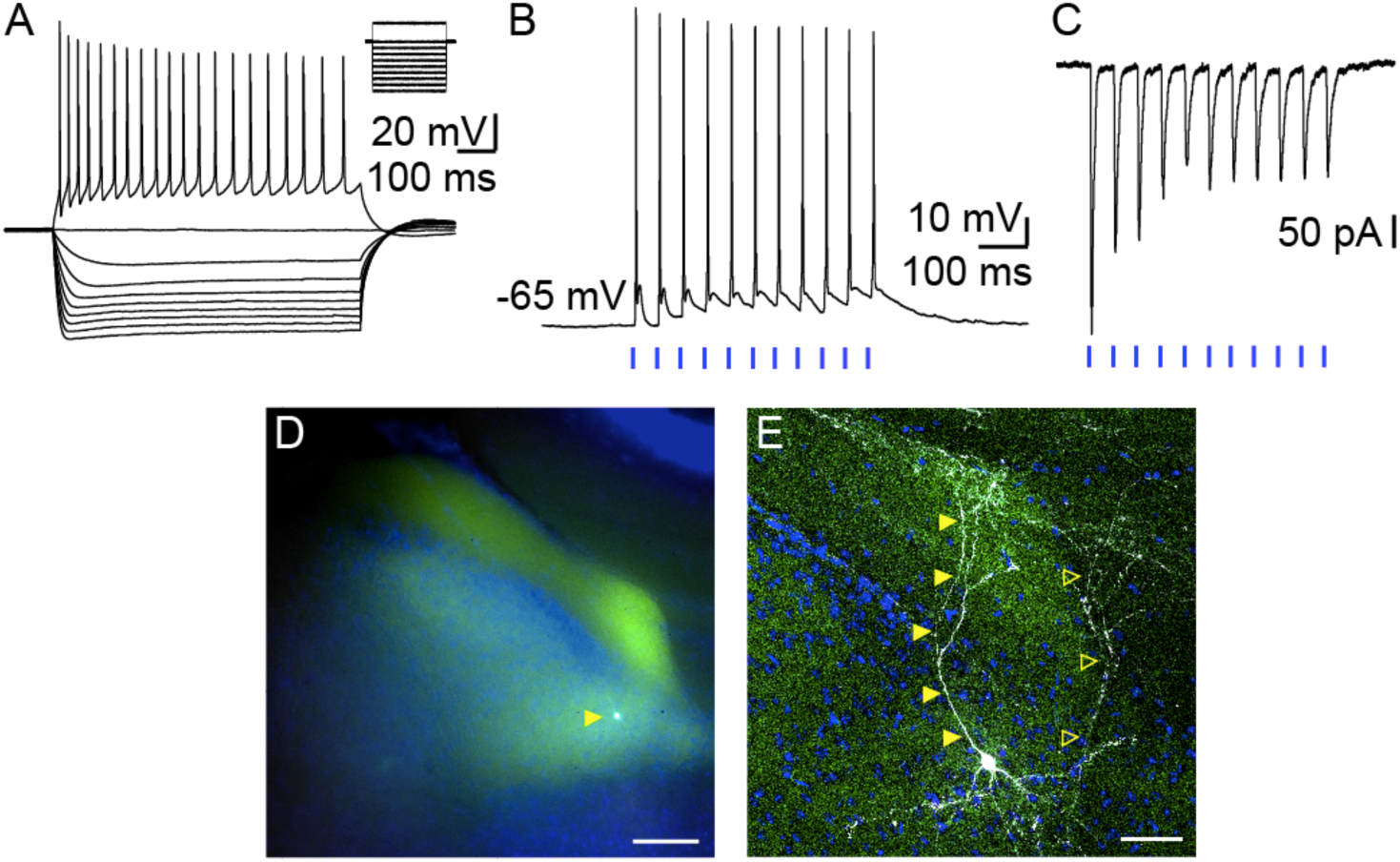
Presubicular layer III neuron: Intrinsic properties, response to light stimulation of thalamic afferents, and posthoc revelation of cell morphology. **A,** Firing pattern and membrane potential variations of layer III neuron for hyperpolarizing and depolarizing current steps. **B, C,** Responses of layer III neuron to 2 ms light stimulations (blue bars) of thalamic axons recorded in current-clamp (**B**) or voltage-clamp (**C**) mode. **D, E,** Layer III pyramidal neuron (white, indicated by a filled yellow triangle) surrounded by thalamic axons expressing Chronos-GFP (green) in presubicular superficial layers with DAPI staining (blue) in horizontal slice imaged with an epifluorescence microscope (**D,** scale bar 100 µm) and a confocal microscope at a high magnification (**E**, scale bar 50 µm). The cell in **A** is indicated with filled yellow triangles. A second, partially filled neuron is present in this slice indicated with empty yellow triangles.
5.10. Record in current- or voltage-clamp postsynaptic responses to whole-field 475 nm LED stimulation of afferent fibers expressing Chronos. Here we stimulate with trains of 10 stimulations of 2 ms duration at 20 Hz (Fig. 2B,C). Light intensity may vary from 0.1 to 2 mW. Response latencies of 2-4 ms are characteristic for a monosynaptic connection.
5.11. To investigate the nature of the synaptic transmission between the long-range afferents and the recorded neuron, different pharmacological agents may be used. Bath application of different receptor blockers allow to define the neurotransmitter that is released and the identity of postsynaptic receptors that are present. To pharmacologically distinguish direct, monosynaptic responses from indirect responses via network activation, add 1 µM TTX and 100 µM 4-AP to the ACSF.
5.12. Wash with original ACSF solution to patch another cell, or transfer the slice containing a biocytin filled neuron in a small vial filled with 4% PFA.
5.13. After an overnight fixation in 4% PFA, wash the slice in 0.1 M PBS (2 × 5 mins, 1 × 20 mins).
5.14. Store in 30% sucrose at 4°C.

### 6. Biocytin revelation

6.1. Transfer fixed slices containing biocytin filled neurons onto a glass blade in a drop of 30% sucrose and perform three cycles of freezing-thawing: place the blade onto dry ice disposed in a Styrofoam box for 1 min until drops of sucrose are completely frozen, then press the blade against the hand palm to thaw.
6.2. Wash the slice three times in 0.1 M PBS (2 × 5 mins, 1 × 1h50), gently agitated. Do not exceed 2h for the last washing.
6.3. Pre-incubate the slice at room temperature for 2h in agitated buffer solution containing 2% milk powder (0.4 g in 20 ml) to saturate non-specific sites and 0.5% Triton X100 (0.1 ml in 20 ml) to permeabilize the membranes, in 0.1 M PBS.
6.4. Incubate overnight at 4°C in a solution containing 2% milk powder, 1% Triton X100, Streptavidin-Cy5 conjugate (1/500) and DAPI (1/1000) in 0.1 M PBS, gently agitated.
6.5. Wash the slice three times in 0.1 M PBS (2 × 5 mins, 1 × 2h). The last wash can last longer, up to 4h, to reduce background staining.
6.6. Before mounting the slice, use an epifluorescence microscope to identify the side of the slice containing the marked cell in a chamber filled with PBS.
6.7. Transfer the slice onto a blade, cell side up, dry it with a paper tissue, mount it using ProLong Gold mounting medium.
6.8. Use an epifluorescence microscope (Nikon, Eclipse TE-2000E) to examine the cell body location or a high-resolution confocal microscope (Zeiss LSM710) for detailed axonal and dendritic morphology (Fig. 2D,E).

## RESULTS

The procedure was used to express a blue light sensitive channelrhodopsin (Chronos) fused to GFP in the antero-dorsal nucleus of the thalamus (ADN), by stereotaxic injection of anterograde Adeno-associated virus. The stereotaxic coordinates were determined according to the Mouse Brain Atlas and tested by injecting 200 nl of fluorescent tracer Fluororuby. The animal was sacrificed 10 min after the injection, and the brain was extracted and fixated overnight. Coronal brain sections were prepared to examine the injection site, which was correctly placed in and limited to ADN (Fig. 1A, B).

In order to express Chronos-GFP in neurons of ADN, we injected 300 nl of AAV5.Syn.Chronos-GFP.WPRE.bGH, and the animal was allowed to recover. Three weeks after the injection, the animal was sacrificed, and acute horizontal brain slices were prepared. Figure 1C shows a brain slice containing the thalamic injection site in the right hemisphere, with GFP expression in green. Upon inspection with an epi-fluorescence microscope equipped with a 4x objective, GFP labeled thalamic axons were observed in the presubiculum (Figure 1C, D). We noted that thalamic axons densely innervated the superficial layers I and III of the presubiculum (Figure 1D).

The activity of presubicular neurons in layer III was recorded in the whole cell patch clamp configuration. Hyperpolarizing and depolarizing current steps were applied while recording the membrane potential variations (Fig. 2A). Data was stored on a PC for later offline analysis of active and passive membrane properties. Presubicular layer III principal cells typically possessed a negative resting potential close to −63 mV and required depolarizing current injections to drive the membrane potential to firing threshold. A full description of their intrinsic properties has been published elsewhere ^10^.

Stimulating ADN axon terminals expressing Chronos-GFP elicited excitatory post-synaptic potentials (EPSPs) in presubicular layer III principal cells in current clamp mode (Fig. 2B). Depending on light intensity, the EPSPs could reach action potential threshold. Postsynaptic responses were also observed in voltage clamp mode as excitatory post-synaptic currents (EPSCs) were elicited (Fig. 2C). Onset latencies of EPSCs evoked by light stimulations were short (median, 1.4 ms; cf. ^9^), indicating a direct synaptic contact between thalamic axons and layer III presubicular neurons. Persisting EPSCs in TTX-4AP condition confirmed this monosynaptic activation (data not shown). It is noteworthy that these cells responded reliably to the light stimulations of afferent axons, with a regular firing pattern.

## DISCUSSION

*In vivo* viral injection to express light sensitive opsins in a defined brain area is a method of choice for the optogenetic analysis of long-range functional connectivity ^9,10,16,17^. Stereotaxic injections offer the possibility to precisely target a specific area of the brain. The co-expression of an opsin together with a fluorescent reporter conveniently allows to evaluate the successful expression and the confirmation of the precise injection site. The use of AAV serotype 2/5 typically restricts expression to the targeted brain region. In this way, a restricted population of neurons becomes transfected, expressing light-sensitive ion channels in their cell bodies and also in their axon terminals. In subsequent *ex vivo* slice experiments it is possible to stimulate these axon terminals with light pulses, directly in their target area, while reading out successful synaptic transmission via patch clamp recording of a post-synaptically connected neuron. The above protocol is robust and convenient, and some additional notes may help successful experiments.

Different types of anesthesia may be used. Here we describe the intraperitoneal injection of a ketamine-xylazine combination as an easy to use, short-term anesthesia with good analgesia^18^. The depth and duration of anesthesia may vary to some extent. Isoflurane anesthesia can be a good alternative to induce faster and to better control the depth of anesthesia. Coordinates of injection sites may be determined with the help of a mouse brain atlas. In practice, coordinates need to be tested, and possibly adjusted if necessary. When positioning the animal in the stereotaxic frame, special attention should be paid to the comfort of the animal, which may also improve the efficiency of anesthesia. The body of the animal should be aligned with the head and the neck. The most critical step in positioning the animal and before craniotomy is the adjustment of the bregma-lambda axis. Especially when targeting deep brain structures, a few micrometers of horizontal deviation will generate errors when lowering the injection needle into the brain. In some cases, one may deliberately choose and calculate an oblique needle trajectory.

The injection volume is a determinant factor for obtaining precisely localized opsin expression. A small volume is ideal to privilege a tightly restricted transfection zone. Higher volumes may be useful to cover the full extent of a large target area. If a large area needs to be covered, such as the septum ^17^, one might choose to place several small injections with a range of neighboring coordinates. The wait time until the *ex vivo* electrophysiological recording is also critical. A minimum time for full expression is necessary, but specificity might decrease if one waits for too long. While 3 weeks seems to be an optimal delay for our experiments ^10^, the necessary delay may vary depending on the virus, its serotype and the distance of the postsynaptic brain region.

The approach described here is even more powerful when combined with injections in transgenic animals. Our previous work has exploited different mouse lines for subtypes of GABAergic neurons, in order to target specifically either PV or SST expressing interneurons for patch clamp recordings ^19^. Simultaneous double recording of neighboring PV and pyramidal neurons or SST and pyramidal neurons then allows to compare the strength of long-range inputs between two neuron types ^10^. This yields results that are standardized with respect to one neuron type, and this standardization is particularly important in case the expression levels of opsins vary between different animals or different slices.

Slice health is of course essential for high quality patch clamp recordings. Constant oxygenation of the slices is crucial, and a slow cutting speed significantly improves slice surface quality. A slice thickness of 300 µm preserves to some extent the microcircuit integrity in horizontal presubicular sections, including the pyramidal neurons with their cell bodies, dendritic and local axonal ramifications and local synaptic connections. The type of light-gated channels chosen to induce activation of afferent fibers will greatly influence the stimulation parameters (duration, light intensity). Chronos is a blue light sensitive channelrhodopsin, and a broad range of illumination wavelengths can be used to activate it (peak sensitivity around 500 nm, even with a minimal light intensity (from 0.05 mW/mm^2^), also activated at 405 nm, and up to 530 nm ^20^). Furthermore, Chronos has fast kinetics properties in comparison to classical ChR2, which enable high frequency stimulations and reliable activation of long-range projections ^21^. In combination with the expression of Chrimson, a red-shifted opsin variant, the independent optical excitation of distinct neural populations becomes feasible.

## ACKNOWLEDGMENTS

We thank Bertrand Mathon, Mérie Nassar, Li-Wen Huang and Jean Simonnet for their help in the development of previous versions of the stereotaxic injection protocol. This work was supported by the French Ministry for Education and Research (L.R., L.S.), Centre National des Etudes Spatiales (M.B.) and Agence Nationale de la Recherche Grant “BURST” (D.F.).

## DISCLOSURES

The authors declare no competing financial interests.

